# Prey capturing and feeding apparatus of dragonfly nymph

**DOI:** 10.1101/536805

**Authors:** Lakshminath Kundanati, Prashant Das, Nicola M. Pugno

## Abstract

Aquatic predatory insects, like the nymphs of a dragonfly, use rapid movements to catch their prey. Dragonfly nymphs are voracious predators that feed on smaller aquatic organisms. In this study, we examine dragonfly nymph (*Libellulidae: Insecta: Odonata)* mouthparts that are used in prey capturing and feeding. In particular, we characterise the morphology of the labium and mechanical properties of the mandibles of the nymph. Additionally, we record and analyse the preying mechanism using high-speed photography. The morphological details suggest that the prey capturing mechanism is a complex grasping mechanism with additional sensory organs that might aid in sensing the surroundings. The times taken for the extension and retraction of labial organ during prey capture was 187±54 ms. The Young’s modulus and hardness of the mandibles samples were 9.1±1.9 GPa and 0.85±0.13 GPa. Gradation in the mechanical properties was also observed in the mandible tip regions with increased properties at the tip end. The overall mechanism with its sensory capabilities provides a unique design to develop bioinspired underwater deployable mechanisms.

## 1.0 INTRODUCTION

Aquatic insects are one of the most diverse groups of animals in small water bodies. While diversity also exists in the broad classification of predators and preys, some of the top predators have been identified as diving beetles, bugs and dragonfly larvae [1]. Aquatic predation presents a complex scenario of mechanics, primarily because of hydrodynamic drag forces experienced during the motion of the predator and its prey-capturing mechanism. Some of these predatory insects use rapid movements to catch the prey. Such rapid movements in many animals is expected to be an outcome of evolutionary pressures such as escaping from predators and catching fast preys [2]. These movements involve jumping and performing predatory strikes, in animals like locust and mantis shrimp, respectively [3,4]. In some of these mechanisms, muscles are used to slowly load spring like structures with potential energy and then quickly release the energy in order to overcome the limitations of temporal contraction of muscles.

Dragonfly nymphs are voracious predators which feed on tadpoles, mosquitoes and other smaller aquatic organisms [5]. They are proved to be a potential bio-control agents of mosquitoes [6]. The predatory strike is performed by using the labium that is modified into a prehensile mask for capturing the fast moving preys [7]. It was shown that the primary mechanisms behind generating jets from the anal valve for self-propulsion and the labial extension for prey capture, follow similar hydraulic dynamics [8]. These hydraulic mechanisms are generated by the use of several muscles on both side of the abdominal diaphragm, thereby generating pressure waves required for propulsion and for labial extension. The mechanics of labial extension of Anax Imperator (*Aeshnidae*) was earlier analysed [9], and the inertial forces and torques acting on the labium were estimated. Earlier studies investigated mechanics of labial extension of other species [9] and suggested a click mechanism at the P1-P2 joint that disengages to release energy [10]. In terms of mechanism, the labial extension occurs by releasing the energy stored using strong contraction of abdomen and thorax muscles [11]. The labial shape of globe skimmer larva (*Pantala flavescens*) used in the present study is different, and has a labial mask that is used to capture the prey. This feature has been reported in previous studies [5], however, the real time dynamics and sensory structures of such a mechanism has not been reported yet.

In the present work, we examine the labium of globe skimmers (*Pantala flavescens*) nymphs that are involved in prey capturing and feeding. To study this, we first record the nymphs underwater using high speed photography. This helps us observe the prey capturing in detail and estimate the time scales involved in protraction and retraction of the labium. Secondly, we measure the mechanical properties of the mandibles of the nymph that are used in feeding the prey after they are captured. Finally, we visualize the mechanism in real time to investigate details of labial extension.. Our study helps to shed further light on the biomechanical aspects of the dragonfly nymph predation.

## 2.0 Experimental materials and methods

### 2.1 Specimen collection and videography

The nymphs of globe skimmers (Insecta: Odonata: Libellulidae) are collected using nets from local, natural and artificial ponds. These nymphs are kept in containers with water collected from the corresponding ponds. The nymphs are then transferred to an acrylic tank of dimension 25 cm (length) × 10 cm (width) × 10 cm (depth) for high speed video recording of the predation process. We use mosquito larvae collected from similar ponds to use as bait for the predator nymphs. Single bait is then laid at some distance from the nymph and the entire process of predation thereafter is recorded using a 1 megapixel high speed camera (Phantom Miro 110).

### 2.2 Microscopy

The images of nymph mouthparts are taken using an optical microscope (Lynx LM-1322, OLYMPUS). Images are captured using a CCD camera (Nikon) attached to the microscope. The dimensions from the images are reported using a standard scale bar in the microscope which checked with a calibration.

SEM imaging is performed directly on the samples which are stored in 100% alcohol and then air dried. Prepared labia are carefully mounted on double-sided carbon tape, stuck on an aluminium stub followed by sputter coating (Manual sputter coater, Agar scientific) with gold. A SEM (EVO 40 XVP, Zeiss, Germany) is used with accelerating voltages between 5 and 20 kV. ImageJ software is used for all dimensional quantification reported in this study (Abràmoff and Magalhães, 2004).

### 2.3 Microindentation and Nanoindentation

Mandible samples separated from the nymphs are embedded in a resin and polished using a series of 400, 800, 1200, 2000 and 4000 grade sand papers. Finally, the sample is polished using a diamond paste of particle sizes in the range of 6 µm and 1 µm, to obtain a surface of minimal roughness. Poisson ratio of 0.31 is used for estimating the modulus [12].

Microindentation experiments (N=2 mandibles from different nymphs) are performed using a standard CSM micro indenter (Vickers) with a load application 50 mN and at a loading rate of 100 mN/min. We use two mandibles from two different nymphs for the microindentation experiments. In nanoindentation experiments (N=2 mandibles from different nymphs), Berkovich indenter is used to perform indentations upto a maximum load of 5 mN at the rate of 1800 mN/min. NanoBlitz3D software is used to map the cross-sectional surface of the tooth samples.

## 3.0 Results and Discussion

### 3.1 Microstructure of the mouthparts

Feeding apparatus of the dragon fly nymph includes several parts such as labrum, mandibles and maxillae. The dissected labium is shown below to show the anatomy of it. It constitutes a postmentum part (P1), prementum part and labial palp (Lp) (Figure 1A and 1B). These parts are moved in co-ordination using concentrated action of muscles, joints and membranous structures [7]. A snapshot taken from the high speed video shows the protraction of the labium with the change in angle of prementum and postmentum of the labium, and opening of labial palp for capturing the prey (Figure 1C).

**Figure 1.**
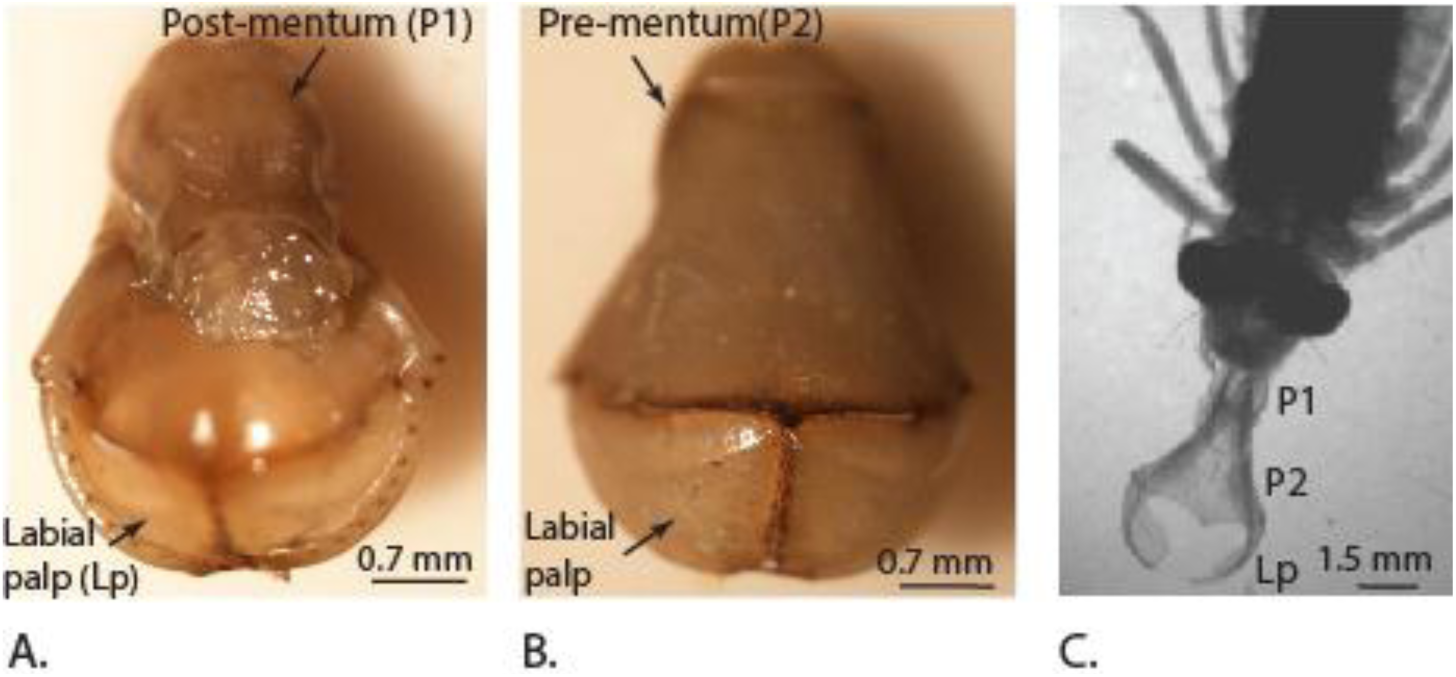
Images showing the labial organ **A.** Dorsal view **B.** Ventral view and **C.** When protracted to capture prey.

The extendable mouthpart labium primary includes prementum, postmentum and the labial palp (Figure 2A). The labial palp consists of two half’s (L and R in Figure 2A) that can open and close, to capture the prey. These two half masks are allowed to move in vertical plane using the corresponding prementum labial mask joints (J1, J2) as shown in Figure 2B. Such joints are also observed in damselfly mouthparts and they were observed to be qualitatively sclerotized using confocal laser scanning method [13]. These joints undergo repeated stress cycles during opening and closure of the masks during life time of the nymph and are resistant to wear. Setae were observed along the rim of the two half masks (Figure 2B). These are hypothesized to be chitin based and help in detecting the either chemical or mechanical signals [13]. Sensillae are observed in a row along the line of contact of the two half masks with the labial palp (Figure 2C). We also observed other type of sensillae that are uniformly distributed on the inner surface of the labial mask (Figure 2D). The sensillae along the edges where the two half’s of the labial mask come in contact, are observed to be in groups of two and three, possibly perform a mechanical function (Figure 2E). They appear very similar to the trichoform sensillum in insects. Such sensillae were observed on the labium of *Crocothemis servilia* [14]. In the antennae of broad-bodied chaser (*Libellula depressa*), coeloconic sensillum was observed and its function is possibly attributed to detecting the temperature variations [15]. We also observed smaller sized sensillae inside sockets (Figure 2E) which appear similar to type ii coeloconic sensillae [16]. The overall macrostructural and microstructural details suggest that the prey capturing mechanism is a complex grasping mechanism with additional sensors that might aid in detecting the surroundings.

**Figure 2.**
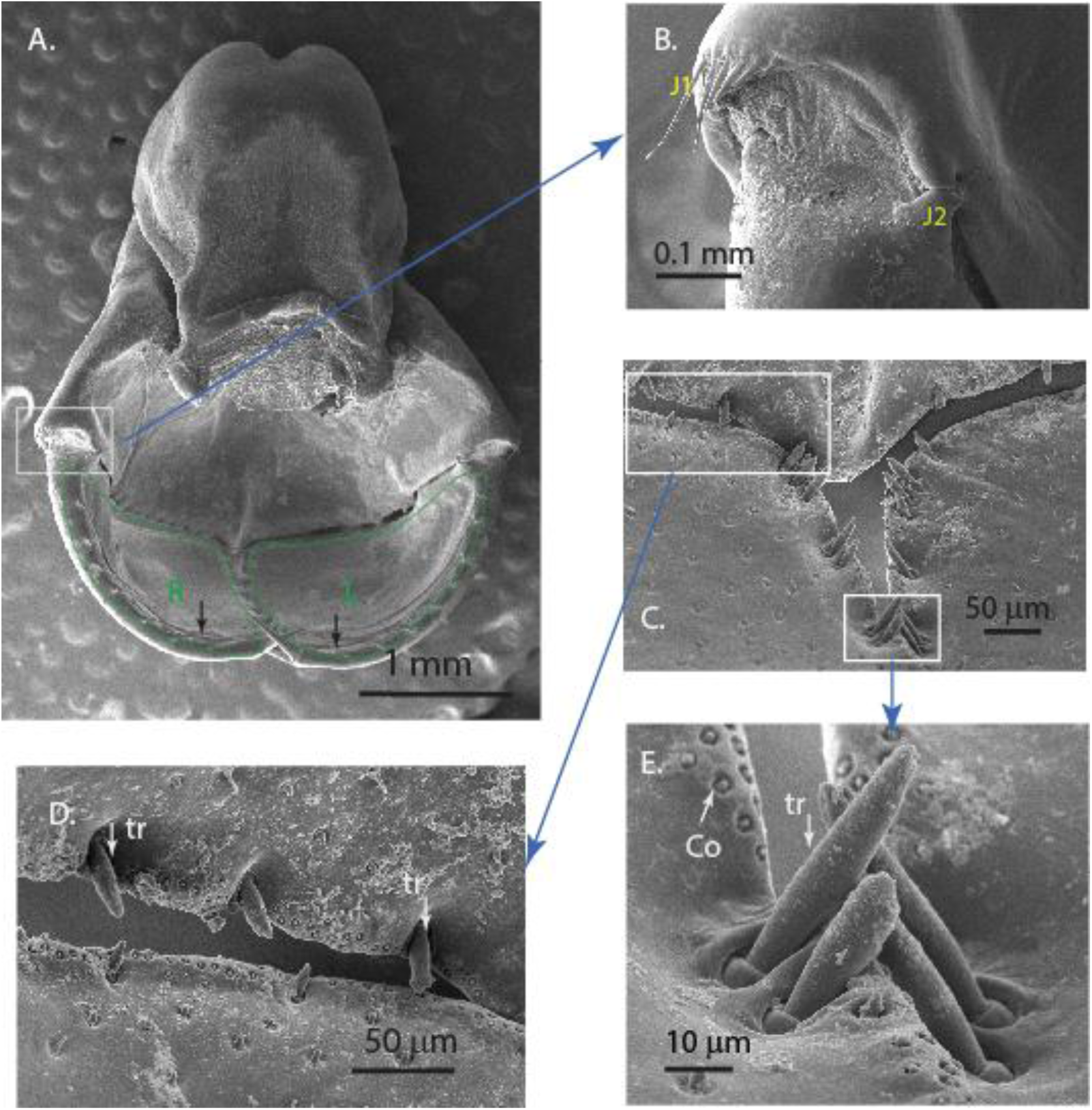
Scanning Electron Micrographs of A) the labium and labial masks (L and R) with bristles (black arrows), B) the joint that plays an important role in the movement of the mouth parts during the prey capture C) sensillae observed on the edges and surface of the labial masks, D) junction of the three mouthparts showing the various types of sensillae E) zoomed region showing the sensillae in pairs along the line of contact of two half labial masks.

### 3.2 Labial movement during prey capture

Our high-speed videos allow us to view the detailed protraction and retraction of labium during the prey capture (Supplementary video 1, video 2). The time taken for full labial extension during the capture of the preys was 63±24 ms (Table 1). This is in close agreement with the time of protraction reported for the Canada darner (*Aeshna Canadensis*) i.e ∼80 ms [11]. The fastest known animal strike by the peacock mantis shrimp (*Odontodactylus scyllarus*) was observed to occur in ∼ 3 ms. Dragon fly nymph predatory strike is not as fast as that of the mantis shrimp but it includes an additional task of opening and closing the labial palps during the time of the strike. The total measured time taken for the extension and retraction of labium during the capture of the prey was 187±54 ms (Table 1). The total time of capture and sweeping, in insects such as the praying mantis (*Tenodera aridifolia sinenis*) was found to be ∼70 ms [17]. The times taken by individual specimens are listed below for more details. The expansion of the prey capturing labium involves kinetic movement of muscles controlled by joints (Figure 3-4).

**Table 1.**
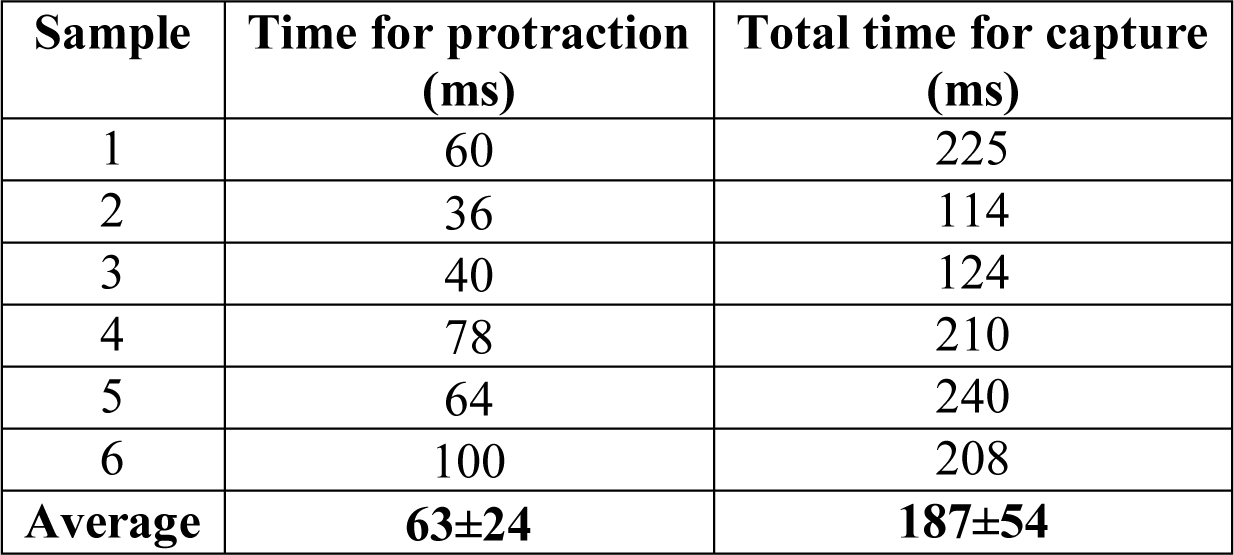
Time taken for protraction of labial palp and capture of the prey.

**Figure 3.**
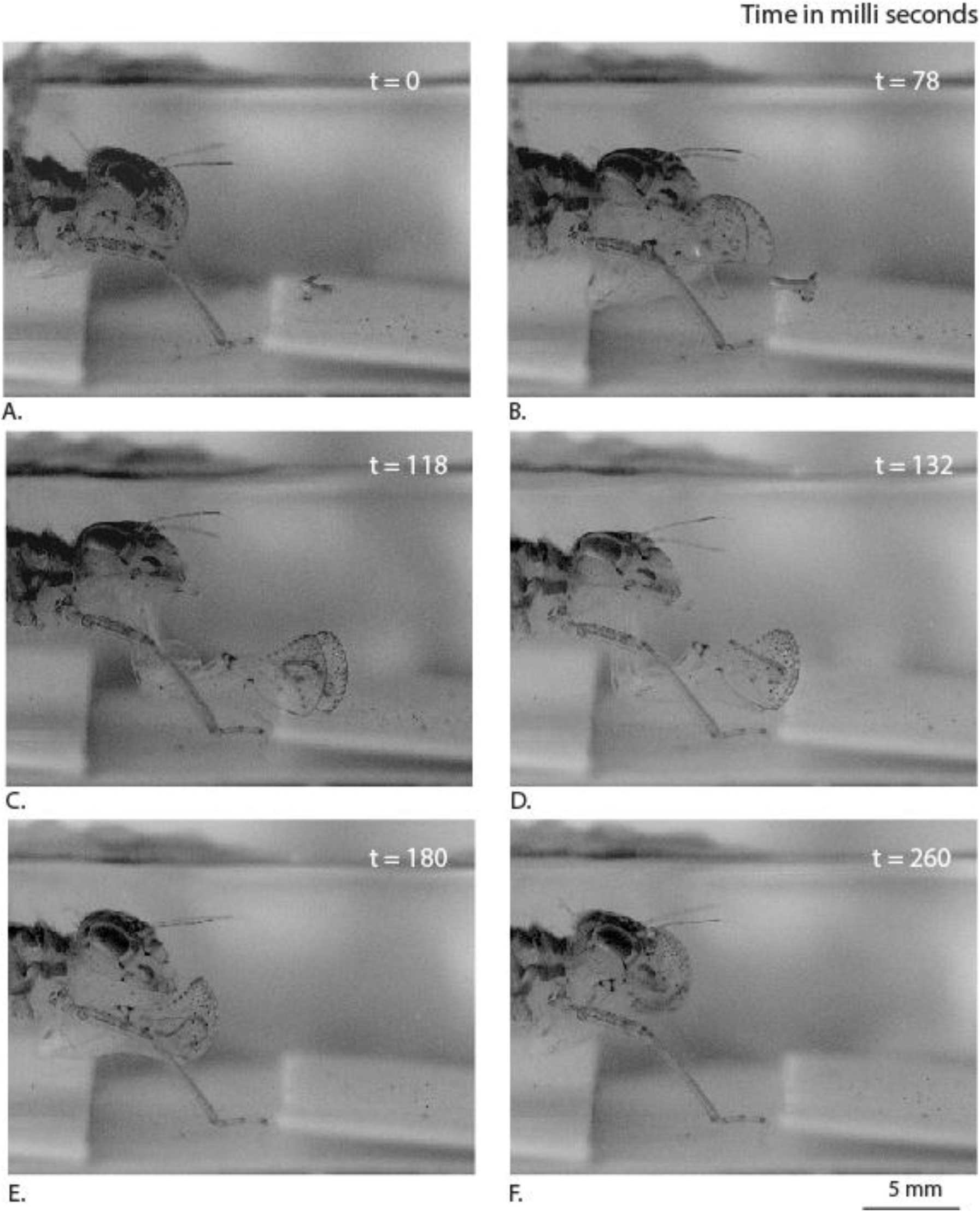
Sequence of movements (side view) used by the dragon fly nymph labium steps to capturing the mosquito larva.

The drag forces acting on the labium during extension can be a combination of viscous drag due to fluid friction, and an ‘added mass’ force. The force due to an ‘added mass’ effect can be understood by noting that the water surrounding the labium is relatively stationary before prey capture, and once the labium opens out, it has to go against the inertial resistance of surrounding water (between panels B and C in Figure 4) in a short duration. This unsteady force can be a significant contributor to the overall drag experienced by the labium. For example, in a study based on swimming scallops, it was shown that the added mass corresponding to transient opening of the shells results in large hydrodynamic forces [18].

**Figure 4.**
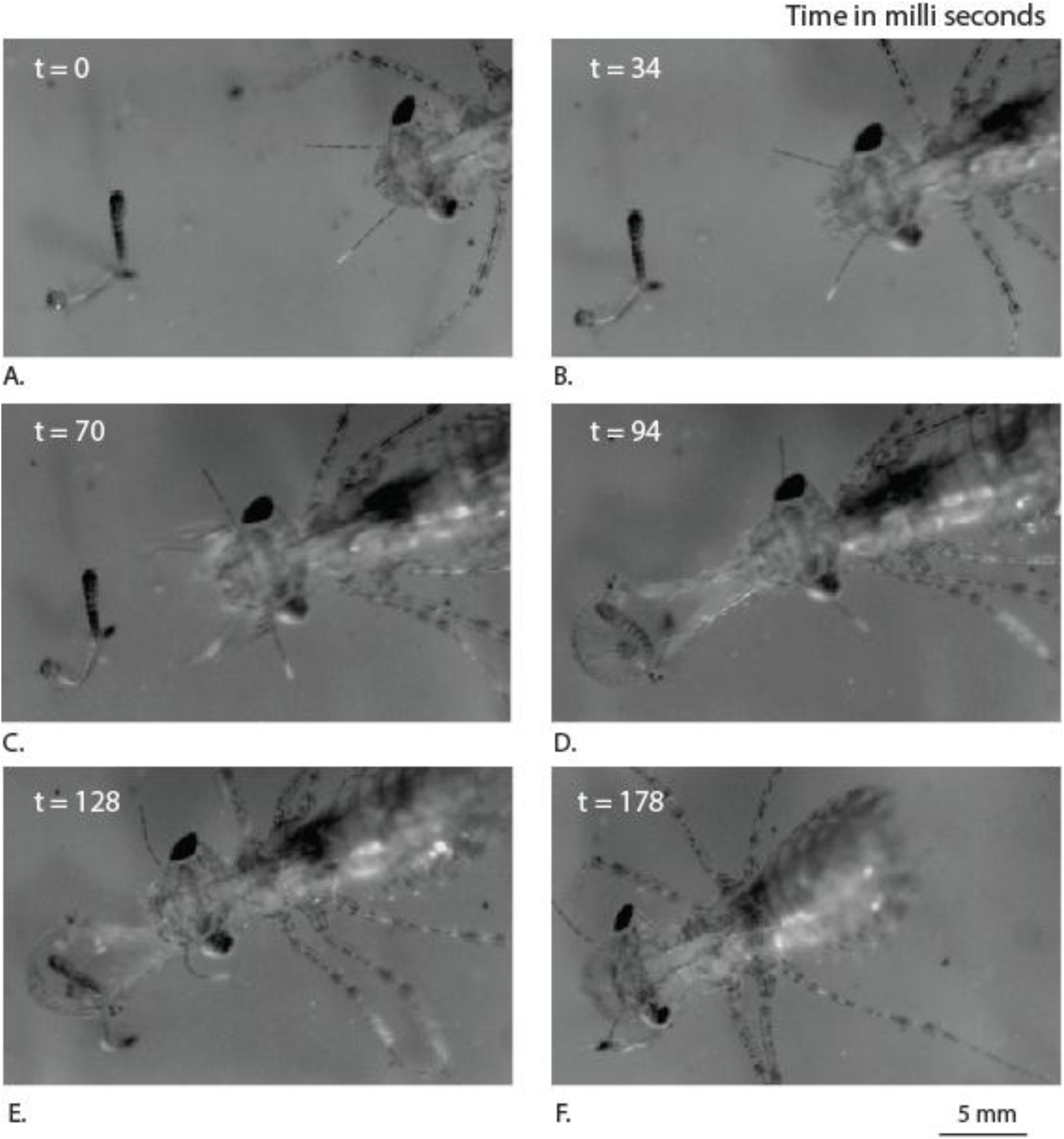
Sequence of movements (top view) used by the dragon fly nymph labium steps to capturing the mosquito larva.

We observe that the labial palps are opened first and the labium is then extended further towards the prey. This is in agreement with earlier observations made on predatory strike of *Aeschna nigroflav* nymph [10]. During this period, when the labium is extending out, viscous drag can be expected to be the dominant force compared to added mass effects, primarily because the labium now slices forward rather than pushing water in the lateral direction. We can clearly see the points (Figure 5: L1, R1) that correspond to the triangular edges of the labial half masks, are rotated vertically upwards to achieve an optimal angle of attack and thereby reduce the viscous drag with the help of each labial mask’s curvature. During this process, the edge of the immovable part of labial mask (Figure 5: B1) also appears to be aligned in such a way to reduce the fluid friction. Thus, the dragon fly nymph appears to hold the labial masks in a way so as to reduce the viscous drag during prey capture and thereby reduce the energy spent. The prementum labial mask joints (J2) (Figure 5) were hypothesized to be strongly sclerotized in a different species of dragonfly nymph [13]. Such joint mechanisms with sclerotized cuticle embedded in flexible cuticle were also observed in the leg joints of frog hoppers [19]. They play an important role in facilitating rotation of the labial palps during opening and closing, multiple times during the life time of the nymph.

**Figure 5.**
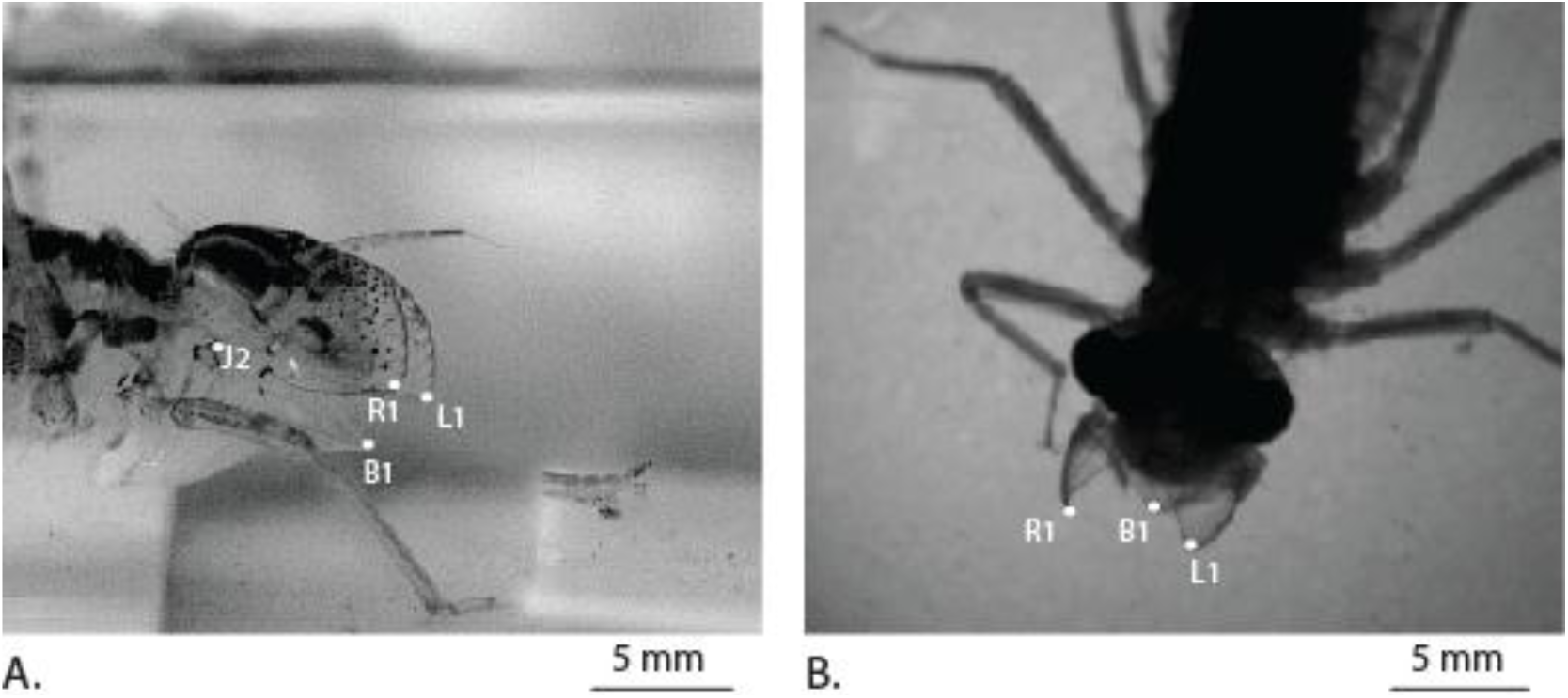
Position of the labial mask just before protraction of the labium A) side view B) top view.

### 3.3 Mechanical properties of the mandibles

Force-depth relation from the microindentation data shows repeatability of the curve (Figure 6A-B). The data analysis shows that the modulus and hardness of the mandibles is 9.1±1.9 GPa and 0.85±0.13 GPa. The mandible cross-section show different regions of sclerotized and non-sclerotized regions (Figure 6C).The indentations from different locations of the mandibles show the Vickers indenter impression (Figure 6 D-E). The measured mechanical properties are similar to that of termite mandibles (Hardness: 0.4-1.2 GPa and Young’s modulus 6-11 GPa) of various species [20] and hardness (0.7-0.9 GPa) of adult leaf cutting ant mandibles [21]. Thus, the mandibles of the nymph appear to be stiff and hard enough to bite through the flesh of the prey.

**Figure 6.**
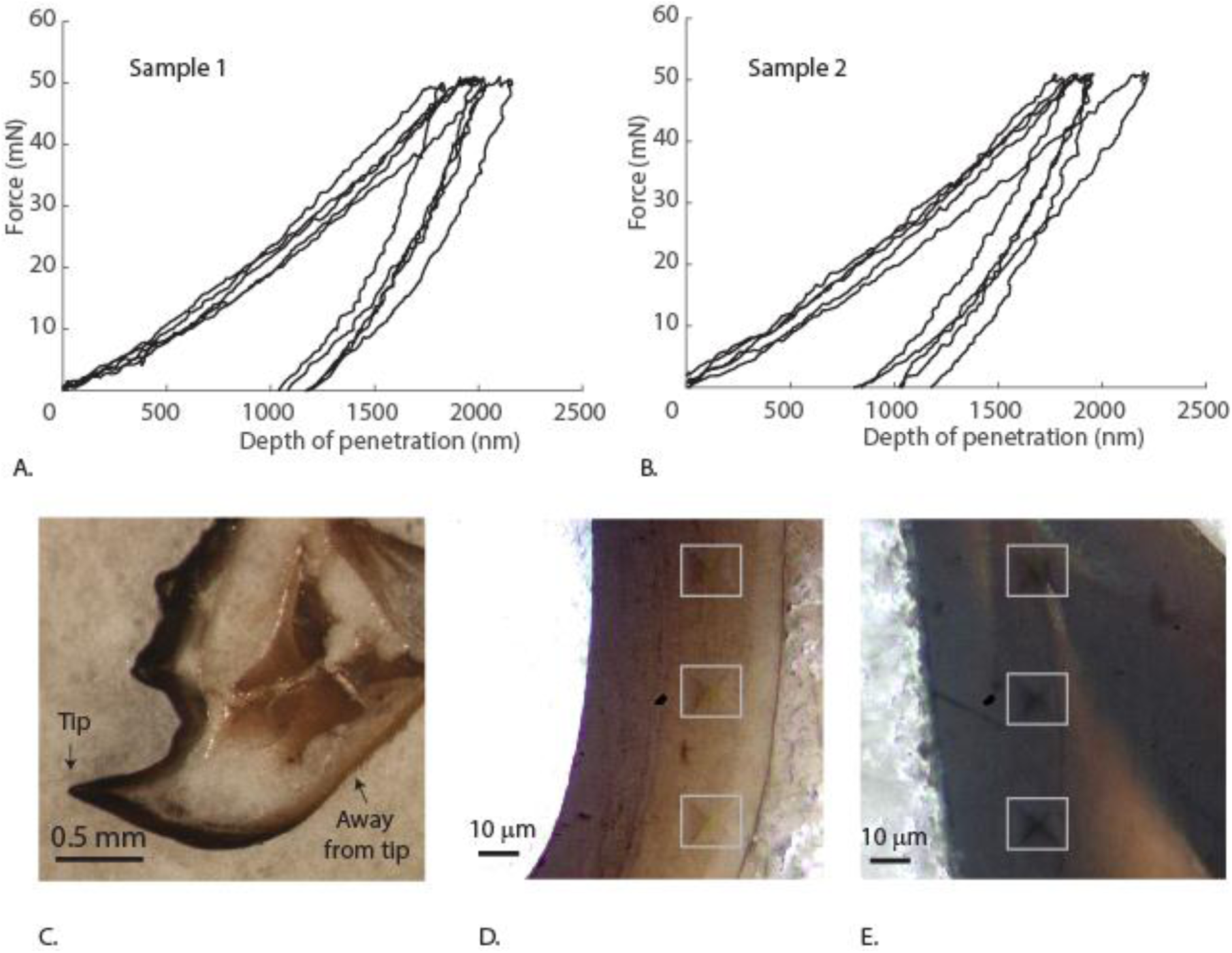
A-B) Force depth curves of microindentation from sample 1 and sample 2. C) Polished cross-section of the mandible. D) Vickers indentation impressions in the away from tip region. E) Vickers indentation impressions in the away from tip region.

Our nanoindentation results show that there is a spread of the data from one region to another, demonstrating the regional differences in mechanical properties (Figure 7-8). The measurements in the region away from the tip (Figure 7) show less variation as compared to the measurements in the tip region (Figure 8). The modulus varies from 6.6 to 8.3 GPa in the base region and, from 2 GPa to 5.5 GPa near the tip region. The hardness varies in the base location from 0.34 GPa to 4.4 GPa (Figure 7D) and, in the tip location from 0.4 GPa to 0.56 GPa (Figure 8D). Most notable, a demarcation in the properties in the mandible tip region is observed in the Young’s modulus and hardness maps (Figure 8A-B). The higher hardness tip regions of the mandibles is known to reduce the wear, as observed in other insect mandible systems [21]. Such gradients were qualitatively observed in the damselfly (*Erythromma najas)* larval mandibles [13]. These gradients can be attributed to the difference in microstructure or metal mediated stiffening resulting in higher properties [22,23]. This needs further investigation to determine the reason behind such variation. The data also suggests that the properties are dependent on the size of the indenter, indicating the role of scaling effects on the mechanical properties.

**Figure 7.**
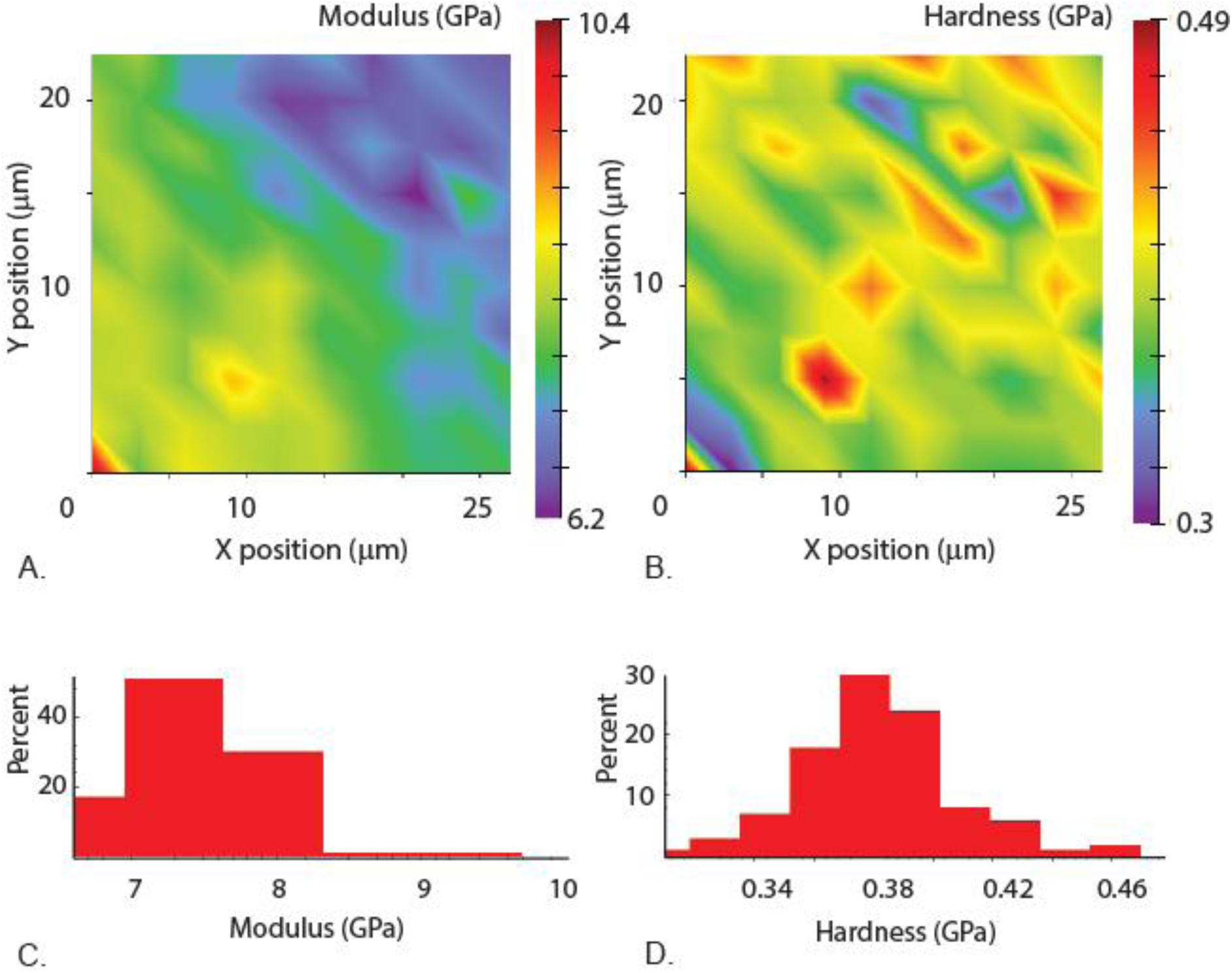
Measurements from the region away from the tip. A) Elastic modulus mapping across the layers. B) Hardness mapping across the layers. C) Corresponding elastic modulus values sorted into bins to show the variation and percentage D) Corresponding hardness values sorted into bins to show the variation and percentages.

**Figure 8.**
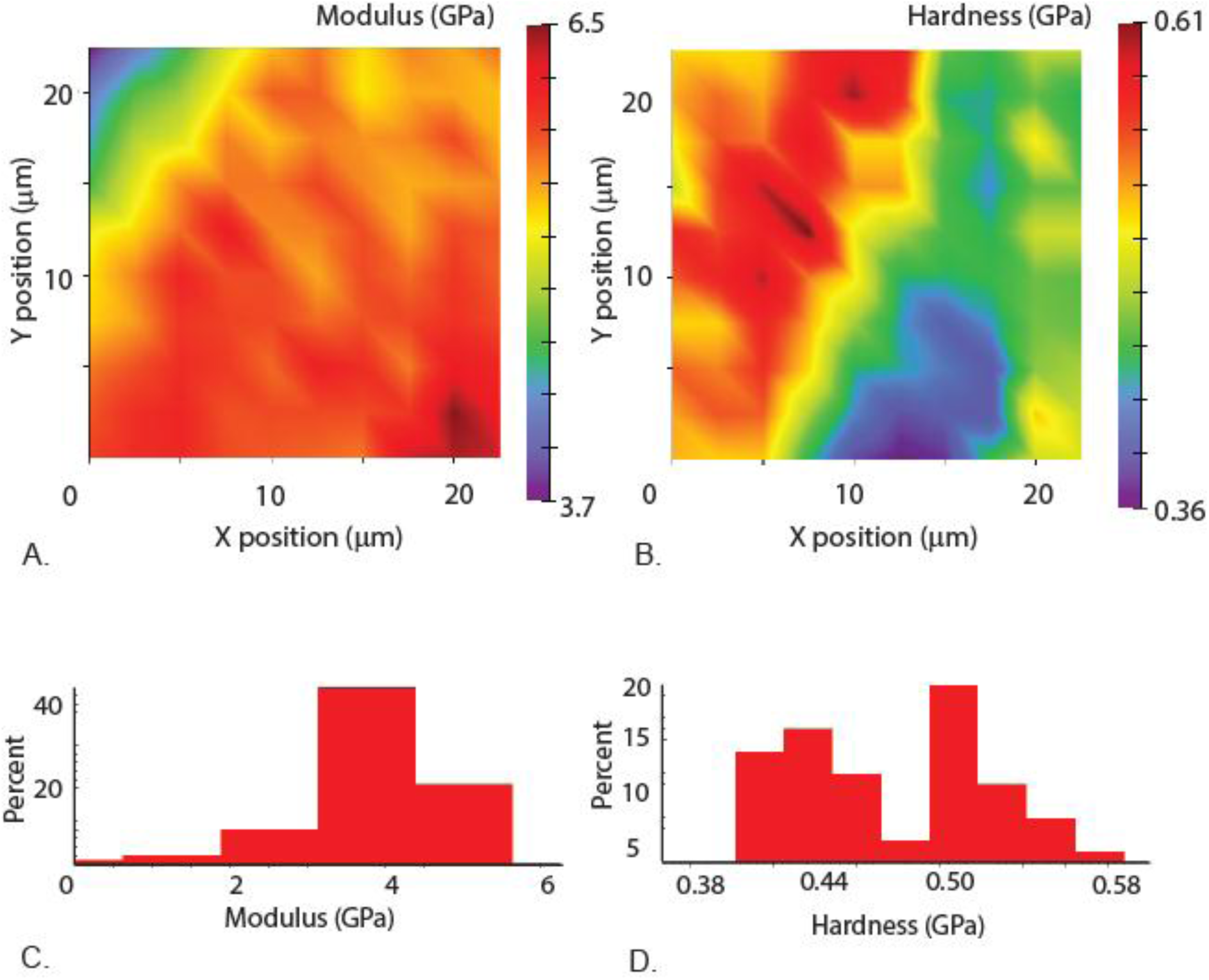
Nanoindentation measurements from the tip region. A) Elastic modulus mapping across the layers. B) Hardness mapping across the layers. C) Corresponding elastic modulus values sorted into bins to show the variation and percentage D) Corresponding hardness values sorted into bins to show the variation and percentages.

## 4.0 CONCLUSIONS

In this study, we discuss the biomechanical aspects involved in prey-capturing mechanism of the dragon fly nymph. We observed that the prey capturing mechanism deploys in a fraction of a second in spite of the inertial and drag forces that are encountered underwater. The nymph mandibles had gradation in mechanical properties as observed in other insect mandibles to reduce wear. The overall mechanism with its sensory capabilities provides an opportunity in the field of bioinspiration for underwater deployable mechanism. Future studies aimed examining the joint morphology can improve the understanding its role in the capturing mechanism.

## ACKNOWLEDGEMENTS

NMP is supported by the European Commission under the Graphene Flagship Core 2 No. 785219 (WP14 “Composites”) and FET Proactive “Neurofibres” Grant No. 732344 as well as by the Italian Ministry of Education, University and Research (MIUR), under the “Departments of Excellence” grant L. 232/2016. We would like thank Manvi Sharma (Centre for Ecological Sciences, IISc, Bangalore) for providing some of the nymph samples and Flow Physics Lab (Dept. of Mechanical Engineering, IISc, Bangalore) for videography.

